# Frequent pulse disturbances influence resistance and resilience in tropical marine microbial communities

**DOI:** 10.1101/2022.07.21.501005

**Authors:** Winona Wijaya, Zahirah Suhaimi, Cherlyn Chua Xin’Er, Rohan Shawn Sunil, Sandra Kolundžija, Ahmad Muzakkir Bin Rohaizat, Norzarifah Binti Md. Azmi, Nur Hazlin Hazrin-Chong, Federico M. Lauro

## Abstract

The Johor Strait separates the island of Singapore from Peninsular Malaysia. A 1-kilometer causeway built in the early 1920s in the middle of the Strait effectively blocks water flowing to/from either side, resulting in low water turnover rates and build-up of nutrients in the inner Strait. We have previously shown that short-term rather than seasonal environmental changes influence microbial community composition in the Johor Strait. Here, we present a temporally-intensive study that uncovers the drivers keeping the microbial populations in check. We sampled the surface water at four sites in the inner Eastern Johor Strait every other day for two months, measured various water quality parameters, and analysed 16S amplicon sequences and flow-cytometry counts. We discovered that microbial community succession revolves around a common stable state resulting from frequent pulse disturbances. Among these, sporadic riverine freshwater input and regular tidal currents influence bottom-up controls including the availability of the limiting nutrient nitrogen and its biological release in readily available forms. From the top-down, marine viruses and predator bacteria (e.g. Saprospiraceae family) limit the proliferation of microbes in the water. Harmful algal blooms, which have been observed historically in these waters, may occur when there are gaps in the top-down and bottom-up controls. This study uncovers the complex interaction between these factors contributing to a low-resistance but high-resilience microbial community and speculate about the rare events that could lead to the occurrence of an algal bloom.

## INTRODUCTION

Microbial ecologists have been fascinated by the impact of disturbances on microbial community succession, resistance, and resilience for several decades. Resistance is the ability of communities to remain stable despite perturbations, whilst resilience is their capacity to return to their original stable state after a disturbance^1, 2^. Many studies, both observational and experimental, have been conducted on disturbance and resilience. However, they do not all produce consistent results, highlighting the complexity of the natural environment.

The city island state Singapore is bordered by two Straits: the Singapore Strait on the southern side separates Singapore from the Riau Islands of Indonesia, while the Johor Strait on the northern side separates Singapore from Peninsular Malaysia. In the 1920s, a 1-kilometer causeway was built on the Johor Strait to connect British Malaya with Singapore, accelerating the area’s economic development. The causeway was built in the middle of the Strait, with years of debris and disrepair preventing water from flowing to either side. As a result, the waters in the inner part of the strait have low turnover rates and nutrient build-up from various anthropogenic inputs^3–7^. The Johor Strait is often at a eutrophic condition, with concentration of nutrients (especially nitrogen) often higher than 10*µ*M for NO_X_ and 40*µ*M for NH_4_, 1 order of magnitude higher than that of the Singapore Strait^8^.

No natural microbial community is immune to changes in environmental conditions, and the microbial communities of the inner Johor Strait are no exception. The Johor Strait is home to many open cage fish farms, and seasonal harmful algal blooms (HABs) have occurred in these waters which had caused billions of dollars in losses to the aquaculture industry^9–14^. However, despite the almost constantly eutrophic conditions, there seems to be some sort of stability in the Johor Strait microbial community as the water does not experience blooms every day. While many observational studies on both bloom and baseline conditions have been conducted in the recent years, none of these studies were done from the perspective of disturbance/resilience of a microbial community despite mentioning some of these disturbances (e.g., sporadic and sudden changes in nutrient inputs and salinity)^3, 5, 6, 10, 12, 13^. Moreover, as the Johor Strait is influence more by short-term environmental changes rather than seasonal ones8, the aforementioned studies were not conducted in a high enough temporal resolution to see the influences of the multitude of pulse disturbances on the microbial community of the Johor Strait.

In this study, we analyse the short-term drivers that affect the Johor Strait microbial populations. We focused on the community compositional resilience, i.e., how the microbial composition remains stable despite disturbances. We sampled water every other day from four sites in the Johor Strait, sequenced the 16S rRNA gene amplicons, and also analysed various water quality parameters and tidal conditions. We posit that, in response to repeated pulse disturbances, the microbial community has evolved to respond to new disturbances with low resistance and high resilience, which might influence how blooms occur in the Johor Strait.

## METHODS

### Data collection

Water sampling and data collection was conducted at the inner Eastern side of the Johor Strait. Sampling sites were chosen due to their accessibility to the Johor Strait waters, either on jetties or land parcels that protrude into the Strait, as shown in **Figure S1** and **Table S1**. Sample and data collection was conducted from 4 November through 28 December 2020, consistently on Mondays, Wednesdays, Fridays, and Saturdays inclusive of these dates (except for samples SNB23 and STL23 on 14 December 2020 due to equipment breakdown).

At each site, surface water samples were collected at 1m depth using a portable pump and filtration system (OSMO)^15^. For DNA extraction, 1.5L of 150*µ*m pre-filtered water were filtered through sterile 0.22*µ*m polyether-sulfone Sterivex filter units (Merck Millipore, Darmstadt, Germany), after which approximately 2mL RNA*later* (Sigma Aldrich, Darmstadt, Germany) was added into the Sterivex, then stored in a dry shipper charged with liquid nitrogen. Duplicate Sterivex samples were collected for DNA extraction. Sample processing took at most 20 minutes between the start of water collection until the addition of RNA*later*.

Temperature and salinity of the water was measured onsite (Extech Instruments, Nashua, New Hampshire, USA). For dissolved inorganic nutrient analysis, 12mL of water from the Sterivex filtrate was collected into acid-washed polypropylene centrifuge tubes, then flash-frozen in liquid nitrogen. Flow cytometry samples were collected from the 150*µ*m pre-filtrate, fixed with electron microscopy grade glutaraldehyde (0.5% v/v final concentration, Sigma Aldrich) at 4°C in the dark before flash-frozen in liquid nitrogen^16, 17^. Chlorophyll samples for two size fractions were obtained: completely unfiltered water, and *<*150*µ*m from the pre-filtrate. For each kind, 50mL of water sample was filtered gently on a 25mm glass fiber filter (GF/F) (Whatman GE Healthcare Life Sciences, Buckinghamshire, UK), then individually wrapped with aluminium foil and flash-frozen in liquid nitrogen.

After each sampling trip, Sterivex filters, cryovials, and chlorophyll GF/Gs were stored at -80°C whilst water samples for nutrient analysis were stored at -20°C until sample processing and analysis.

#### Tide and rain data acquisition

Half-hourly tide measurements were acquired from the Maritime and Port Authority of Singapore (https://www.mpa.gov.sg/) from the government-run Sembawang Tide station (1.465° N, 103.835° E) (**Table S3**). Current strength and direction were calculated by the slope of the tidal data, i.e., the change in tide height over a unit of time. Dates of peak neap and spring tides were estimated from moon phases, which were then used to calculate the number of days between sampling and its nearest peak of neap/spring tide (**Table S2**). Rain data was acquired from the Meteorological Service Singapore (http://www.weather.gov.sg/climate-historical-daily/) using Sembawang Station (1.4252° N, 103.8202° E) for eastern sampling sites and Admiralty Station (1.4439° N, 103.7854° E) for western sampling sites.

### Sample processing

#### Dissolved inorganic nutrient concentration

Samples for dissolved inorganic macronutrients were thawed at room temperature and immediately measured on a SEAL AA3 High-Resolution AutoAnalyser (SEAL Analytical, Norderstedt, Germany). The nutrients measured were phosphate (PO_4_), silicate (Si), ammonia (NH_4_), nitrite (NO_2_), and NO_x_, all in *µ*mol/L. nitrate (NO_3_) was calculated as NO_x_ -NO_2_, while total Dissolved Inorganic Nitrogen (DIN) was calculated as NO_x_ + NH_4_.

#### Chlorophyll concentration

GF/F filter of each sample was placed into acid-washed centrifuge tubes containing 90% molecular-grade acetone and incubated in the dark at 4°C overnight. Samples were then centrifuged at 500G for 10 minutes. The supernatant (2mL) was then aliquoted into a disposable cuvette for measurement using the FluoroMax spectrophotometer (Horiba).

#### Flow cytometry (FCM)

Our FCM analysis protocol was optimised for counting viruses based on Brussaard (2004)^16^ and Brusaard et. al (2010)^17^. Briefly, gluteraldehyde-fixed samples were diluted using Tris-EDTA buffer and stained using SYBR Green I (Invitrogen, Life Technologies, Eugene, Oregon, USA). Samples were run at the slow flow rate (around 10*µ*L/s) on the CytoFLEX benchtop flow cytometer (Beckman Coulter, Brea, California, USA) equipped with blue (488nm) and violet (405nm) lasers specifically to distinguish virus particles. Virus and bacterial populations were gated using the CytExpert software (Beckman Coulter), validated against fluorescent microscopy counts and serially diluted positive controls. The raw FCM data was then processed further in the R statistical environment^18^.

#### DNA extraction, amplification, sequencing, and data processing

DNA samples were extracted in random order to minimise bias and batch effects. We followed the default factory protocols of the DNEasy Powersoil Pro kit (QIAGEN, Germantown, Maryland, USA) with a few modifications. Firstly, filter units were rinsed with 1X phosphate-buffered saline to remove the RNA*later*. The Sterivex casing was broken to expose the filter inside, which was inserted into Powersoil Pro Bead tubes (prepared as per the first step of the default protocols), then incubated at 70°C, 500rpm for 15 minutes, twice. Molecular grade Phenol-Chloroform-Isoamyl Alcohol (25:24:1 v/v) (Sigma Aldrich) was added into the bead tubes, after which the samples were subjected to rest of the factory protocol starting at the 10-minute vortexing. DNA was eluted into 60*µ*L of nuclease-free water and quantified using a Qubit 2.0 fluorometer (Invitrogen, Life Technologies).

Of each sample, PCR was done in triplicates using KAPA HiFi HotStart ReadyMix (Roche, Cape Town, South Africa) with primers that have the standard Nextera Illumina adapter attached at their 5’ end. The primer pair, 926WF (5’-AAA-CTY-AAA-KGA-ATT-GRC-GG-3’) and 1392R (5’-ACG-GGC-GGT-GTG-TRC-3’)^19^ was used specifically because it targets the V6-V8 hypervariable region of the SSU rRNA gene of all three bacteria, archaea, and eukarya at a considerably high coverage for environmental samples^20, 21^. After 22 cycles of amplification, triplicate amplicons of the same samples were pooled and then cleaned using KAPA HyperPure Beads (Roche).

Amplicon library preparation and sequencing was conducted at the sequencing facility at Macrogen APAC (South Korea). Briefly, a second round of PCR was done to attach dual barcodes to each sample for multiplexing. The pooled library was then sequenced on an Illumina MiSeq machine. The demultiplexed .fastq files were returned to us and further processed in the R statistical environment^18^.

Adapter and primer sequences were removed from the reads using cutadapt^22^. The reads were then subjected to processing using the DADA2 algorithm^23^. The Silva rRNA database version 132^24^ was used to classify the taxonomy of each amplicon sequencing variant (ASV), after which the PR2 database^25^ was used to re-classify the eukaryote ASVs. Afterwards, chlorophyll and arthropod sequences, as well as ASVs with less than 10 reads across samples were removed. Chlorophyll sequences were removed to avoid double counting photosynthetic organisms, whilst arthropods are usually bigger than 150*µ*m and thus the DNA detected most likely have come from their body parts or eggs, and too-rare ASVs might be an artifact of de-noising or wrongly read sequences. ASV abundances between technical duplicates were normalised and then averaged. As shown in **Figure S2**, the Bray-Curtis distance between different locations of the same day is generally larger than the distance between technical replicates.

### Statistical Analysis

The R package *vegan*^26^ was used in most statistical analyses of the processed sequencing data: calculation of alpha diversity indices (Shannon & Simpson indices), pairwise community dissimilarity using the Bray-Curtis distance, dimensionality reduction of these distances using non-metric multidimensional scaling (nMDS), analysis of environmental parameters using *envfit*, as well as Canonical Correspondence Analysis (CCA). The packages *phyloseq*^27^ and *ggplot2*^28^ were also used to visualise the results of these analyses, and *ggpubr*^29^ was used for statistics on graphs. Co-occurence networks were built using the *SpiecEasi* algorithm^30^ and analysed using *igraph*^31^, all in the R environment. Networks were then visualised and exported using Gephi^32^.

Principal Components Analysis (PCA) was conducted using the *princomp* function, while correlation analysis between environmental variables were conducted using the *cor* function. Visualisation of correlations were done using *corrplot*^33^.

Distance-based phylogenetic trees were made to classify unclassified Saprospiraceae ASVs. In-silico PCR (https://github.com/egonozer/in_silico_pcr) was conducted on genomes downloaded from NCBI Refseq^34^ with primers as the above primer pair. Alignment was calculated using MAFFT^35^ and trees were constructed using FastTree^36^, then visualised using FigTree. *E. coli* strain KS-12 was used as an outgroup to root the phylogenetic tree.

## RESULTS

### General overview

During the course of sampling, the average salinity and temperature of the East Johor Strait waters were 30.1psu and 25.3°C, respectively. Approximately 35 rain days were recorded during the duration of sampling; the maximum number of days without rain was 4 days (29 Nov – 2 Dec 2020, inclusive). This trend is consistent with historical rain reports of Singapore (http://www.weather.gov.sg/climate-climate-of-singapore/).

Despite receiving approximately the same amount of rain as the western sites, the site closest to the mouth of the Strait (SBW) had a significantly higher salinity (**Figure S3**). The western sites seem to be more affected by tidal mixing, with a higher community dissimilarity closer to neap tides when tidal mixing was low (**Figure S4**). Otherwise, there were no consistent differences in the nutrient concentration and other observed environmental parameters between the sites. While the concentrations of dissolved nutrients did vary throughout the time series, variations were consistent across the four sites (**Figure S5**).

### Amplicon Sequencing Results

A total of 9,543,682 sequencing reads were recovered, with 2,847 ASVs across 118 samples after trimming and a mean of 31,868 reads per sample. Bacteria of the phyla Bacteroidetes, Cyanobacteria, and Proteobacteria make up the majority of all organisms captured; the *Roseobacter* HIMB11 (ASV0003) and cyanobacteria PCC 6307 (ASV0004 & ASV0007) were the top three most abundant ASVs in the Johor Strait. Some past HAB taxa, as listed in previous studies conducted in the region^5, 9, 12–14, 37^, were observed in our samples: dinoflagellates of genus *Chaetoceros* and family Kareniaceae, as well as diatoms of genus *Thalassiosira*. These taxa were only found in very low amounts at a mean of 82 reads per sample, however, they were present in all but 37 samples in our time series.

### Correlations

Correlation analyses were conducted between environmental parameters, as shown in **Figure 1**. A notable correlation interaction can be observed between the tides, chlorophyll and nitrogen ratios (DIN:P and DIN:Si). There tended to be higher chlorophyll concentrations during spring tides when tidal-driven mixing was the highest. Vice versa, a lower chlorophyll concentration was observed during the neap tide. Interestingly, chlorophyll concentrations were positively correlated with the N ratios but not with the raw N values. The rain also brought higher N ratios, which was unexpectedly not followed by the increase in chlorophyll concentrations.

**Figure 1.**
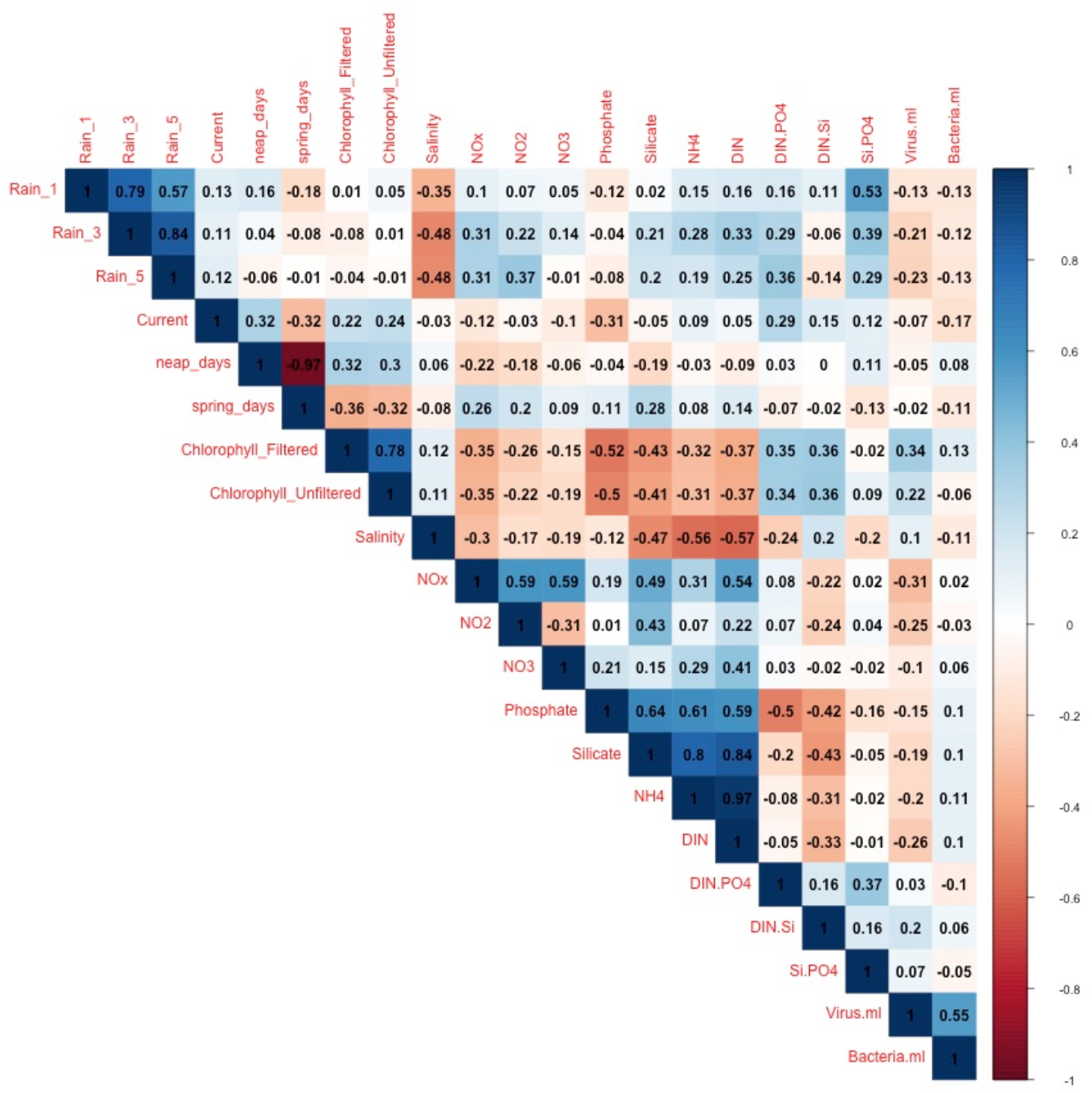
Correlation of various environmental variables measured for all sites

Across the domains, virus counts showed a positive correlation with bacteria, in both the FCM gated counts and chlorophyll concentrations: high virus counts were found when high bacterial counts were also observed, and vice versa. This is further seen in **Figure 3** where the virus and bacterial FCM counts from nearly all points were in sync with each other in their increases and decreases.

**Figure 2.**
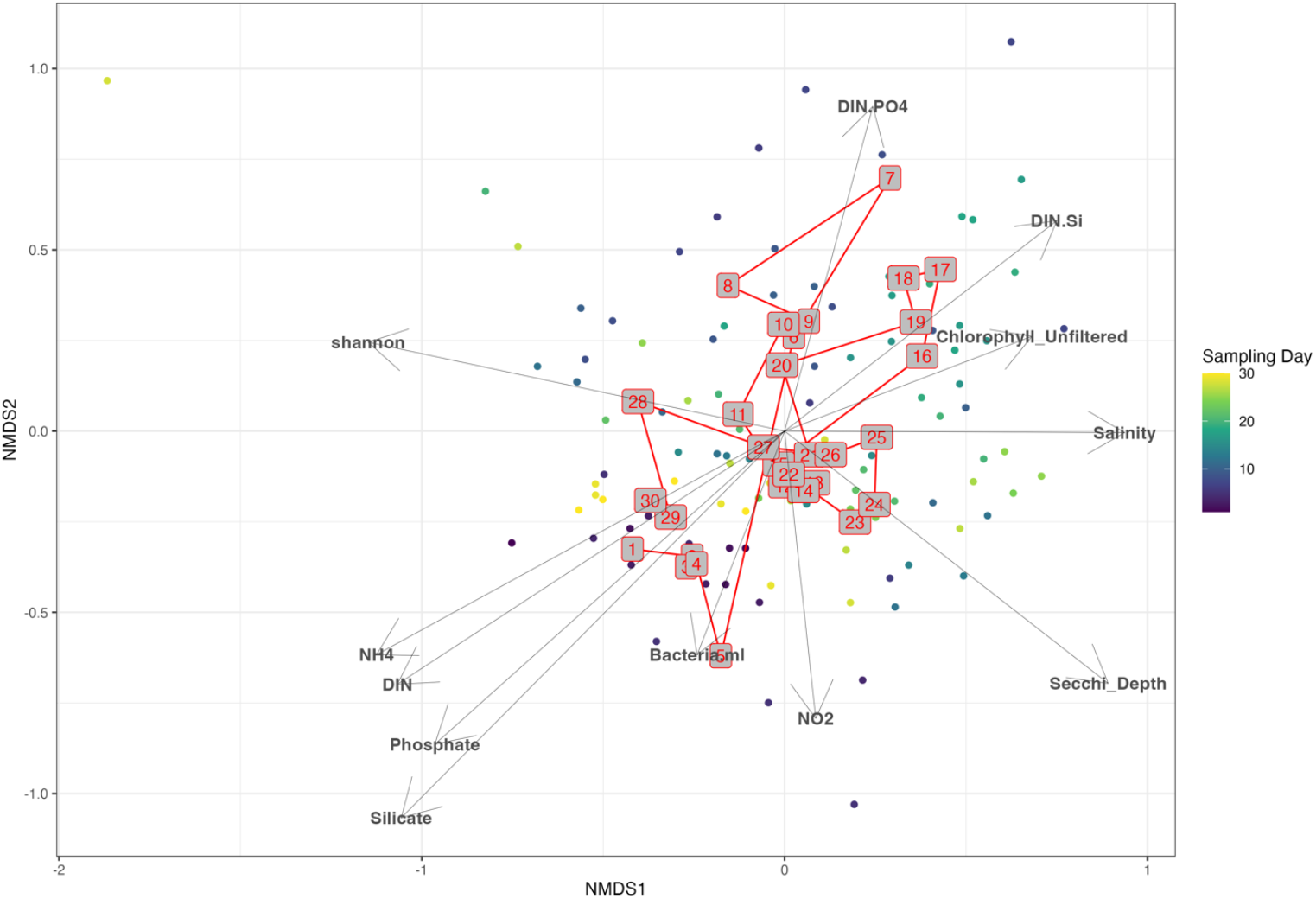
Non-Metric Multidimensional Scaling (nMDS) on the Bray-Curtis distance between each sample. Arrows show *envfit* results, while the red numbers in grey boxes show the centroids of each sampling day.

**Figure 3.**
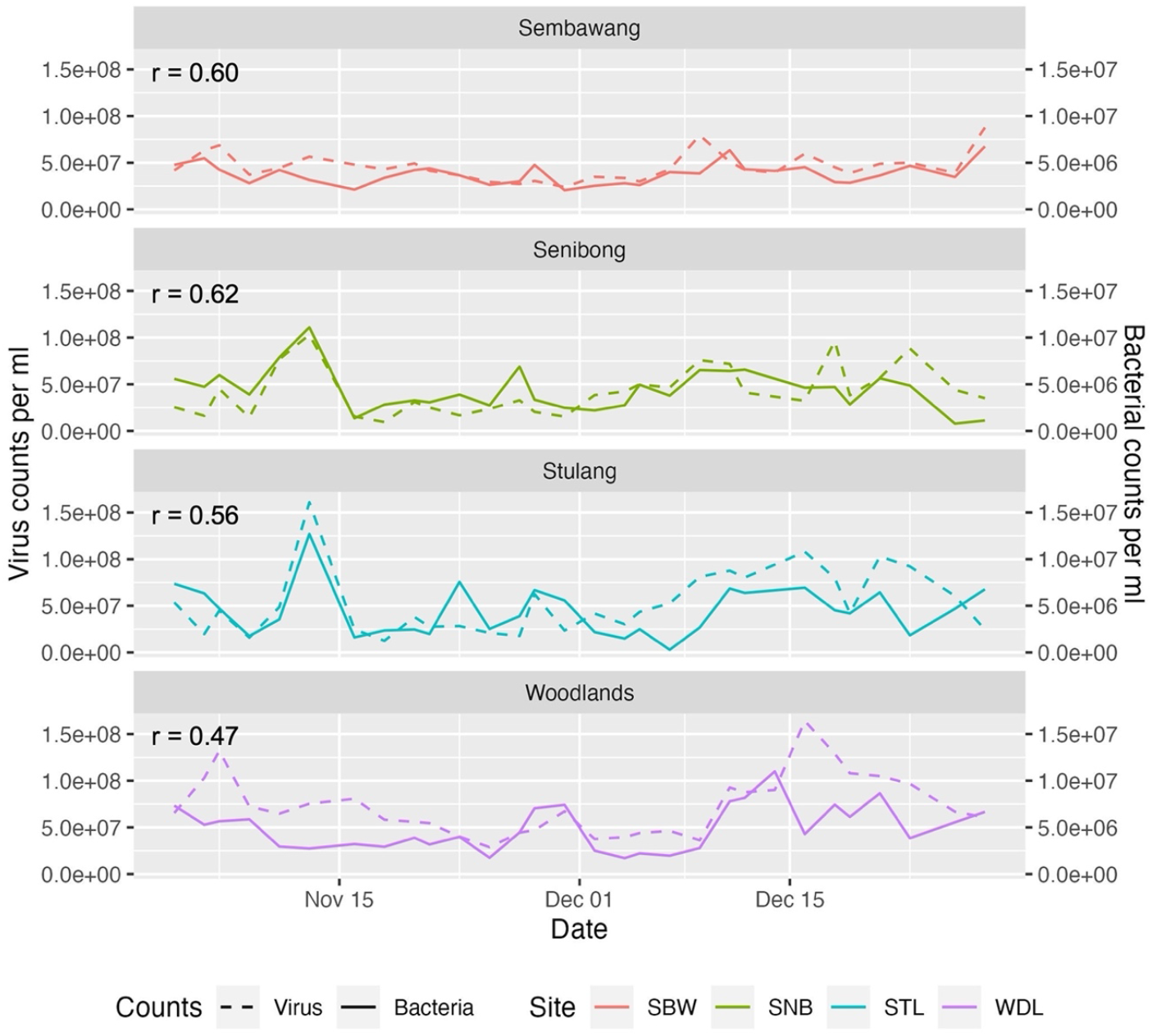
FCM counts of bacteria & virus particles, showing the high correlation between bacteria and its top-down controller viruses

### Trajectory

Dimensionality reduction of the Bray-Curtis distance using non-metric multidimensional scaling (nMDS, **Figure 2**) which shows the community succession between one sampling day and the next seemed to revolve around a common stable state, circling the midpoint of (0,0). Furthermore, *envfit* analysis revealed many water quality parameters that were highly correlated with the samples. No significant autocorrelation of environmental parameters were observed (**Figure S6**).

Principal components analysis (PCA) returned the variables that contributed heavily to the variation in the data in attempt to reduce its dimensionality. The first axis represents 34% of the variation found in the data, which was mostly contributed by two ASVs related to *Cyanobium* PCC 6307 (89.3%). The second axis represents 21% of the variation, which is mostly explained by changes in the abundance of the Rhodobacteraceae strain of HIMB11 (68.6%). These two taxa represented the organisms that explained most of the variation in our time series, which coincided with being the top three most abundant ASVs in our data.

Canonical correspondence analysis (CCA) showed how each ASV might have been influenced by nutrients as well as the salinity of the water (**Figure S7**), with **Table S4** summarising the sensitivity of a few notable taxa. To illustrate this further, **Figure 4** shows a time series of these taxa and the relative concentrations of nutrients for comparison. No taxa seemed to take over the community for an extended period of time; as the abundance of one species rose, others fell, and vice versa. For example, the small peak of HIMB11 after the 1st of December was characterised by the lower abundance of other taxa, and coincided with the relatively high DIN:Si, Salinity, and DIN:PO4.

**Figure 4.**
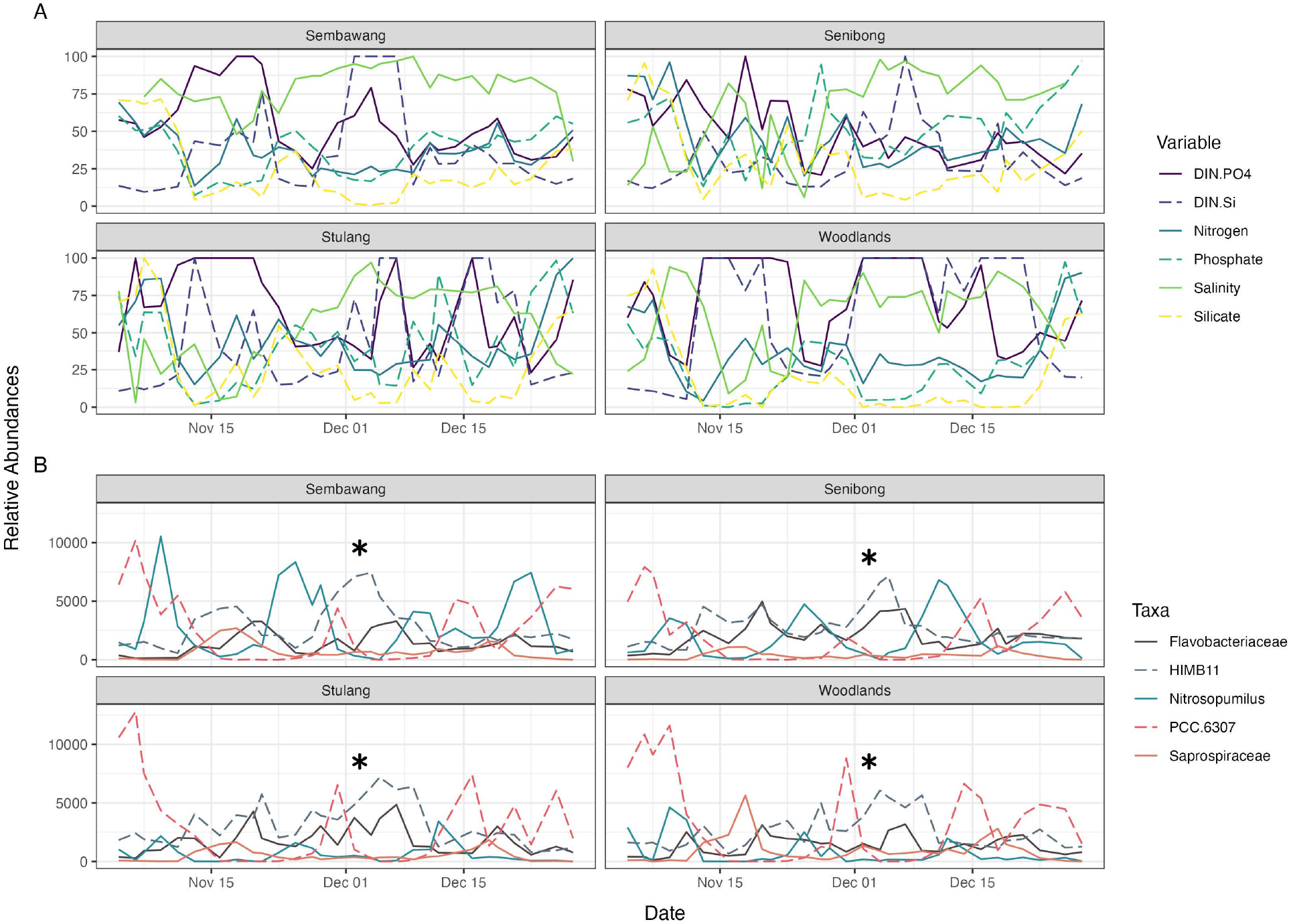
Time-series of nutrients and a few of the most abundant taxa, note the changing peaks of the nutrient availability and dominant taxa. (A) Nutrient data have been normalised between 0 to 100 to set everything on the same scale and order of magnitude for easier comparison. (B) Amplicon reads have been normalised to the median total read count. The asterisks (*) note down the position of the HIMB11 peak mentioned in the main text.

### Network Analysis

Co-occurence networks show potential interactions between taxa, such as predatory (-), parasitism (+), symbiosis (+), and antimicrobial (-) relationships. However, networks could not differentiate between the different types of interactions; it does not make a distinction between parasitism and symbiosis as both are positive relationships. Furthermore, it is very difficult to experimentally confirm these relationships, and thus co-occurence networks are more useful as a hypothesis generator rather than to be used as confirmatory analysis^38–40^.

Despite its many limitations, one interesting point can be noted from the results of our co-occurrence network in **Figure 5**. The family Saprospiraceae (especially ASV0037) has an Eigenvector Centrality of 1, meaning that its node is well connected to other well-connected nodes, making it a very influential node. Therefore, Saprospiraceae is a potential ‘keystone taxa’ in the Johor Strait microbiome, that is, it has the potential to have a crucial role in microbiome structuring in the Johor Strait. Keystone taxa do not need to be the most abundant species in the community; there are instances whereby the rare microbial taxa in the microbiome are disproportionately important for the well-functioning of the entire system^41^.

**Figure 5.**
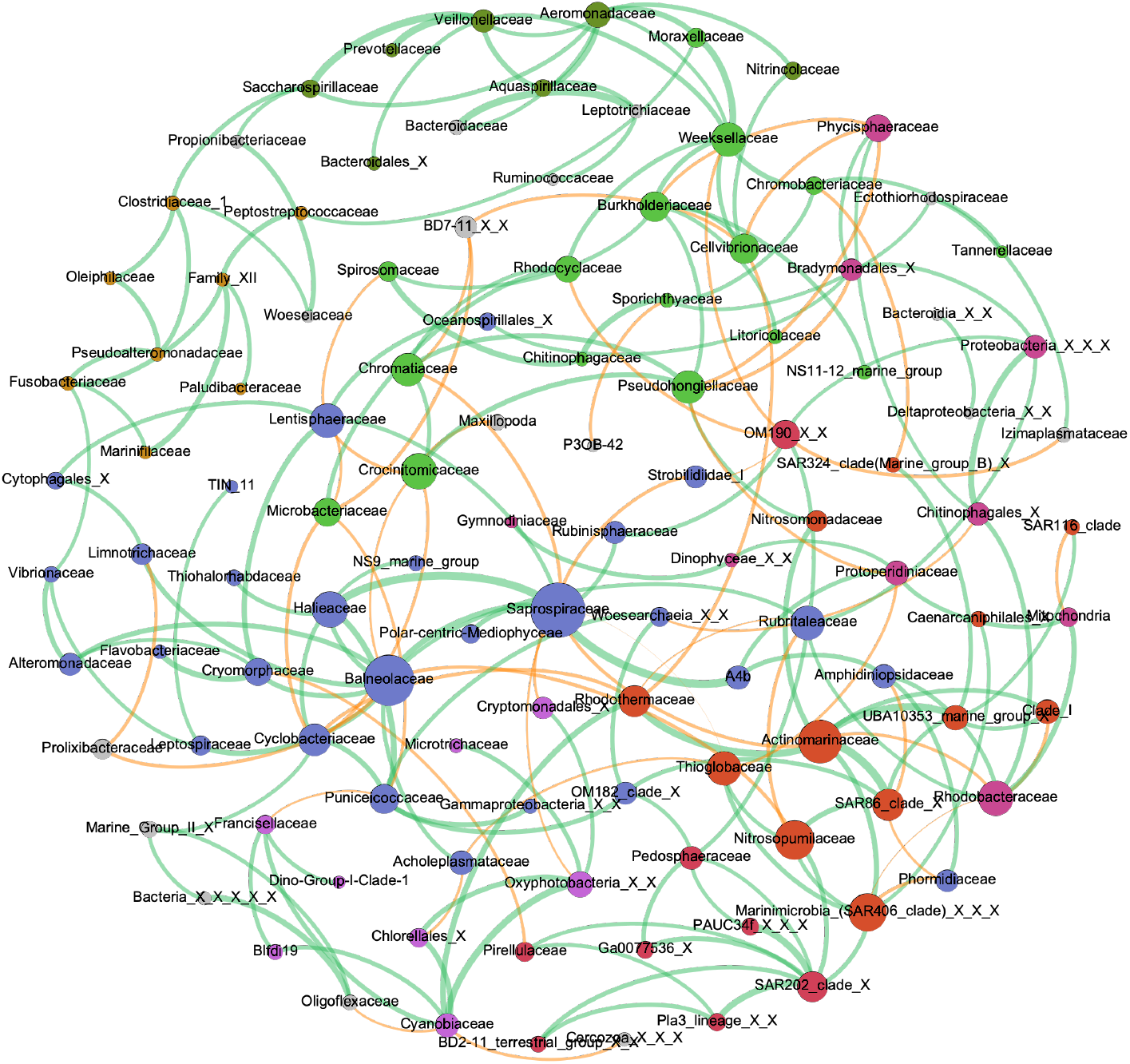
Co-occurrence network analysis at the Family level, showing Saprospiraceae as the potential keystone taxa. Green edges show positive co-occurrence, while orange edges show negative co-occurrence. Colours of the nodes show the potential clusters that co-occur together.

As shown in **Figure S8**, the phylogenetic tree developed from extracted Saprospiraceae 16S rRNA genes in the NCBI RefSeq database show that the most abundant Saprospiraceae-related ASV (ASV0037) was found to be most closely related to *Aureispira maritima*, a motile Saprospiraceae first isolated from barnacle debris in Thailand^42^.

## DISCUSSION

### Stability of Johor Strait microbial community

During the duration of the experiment, the community structure remains in a relatively stable state, with no single taxa becoming dominant at any point during the sampling period. Resilience was observed despite the frequent perturbations experienced in the area (i.e., irregular nutrient-enriched freshwater pulses from various sources, the temporally consistent saline seawater intrusion that comes with the tidal currents, and immigration of foreign populations in runoff water, among other things) (**Figure 2**). Each of these disturbances resulted in a now somewhat-predictable change in the community composition.

As this has been occurring long before sampling, we propose that repeated disturbances may have driven selection, adaptation, and/or diversification of the community. As a result, the microbial community that we observe today is resilient and is able to tolerate these frequent disruptions. A history of repeated disturbances may promote community resilience through these methods, according to earlier studies and evaluations.^1, 2, 43, 44^

**Figure 4**. shows how the most common taxa in the Johor Strait alternately became the most abundant species, but none of them ever dominates the community for an extended period of time. The changing environmental conditions constantly open up new niches which can be filled better by other taxa. During the periods when they are not the most abundant species, they are still present in small amounts until conditions no longer favour the current dominant taxon and, instead, favour a particular taxon’s growth the most. The changing peaks in **Figure 4** are mostly made up of organisms related to HIMB11, *Nitrosopumilus*, PCC6307, and on a few instances, Saprospiraceae. These taxa would need rapid growth rates to be able to keep up with the ever-changing environmental conditions and nutrient availability. In short, **Figure 4** shows the slightly different niches rapidly opening up and are immediately filled in by the taxa that fits them best and fastest.

These repeated pulse disturbances observed may have also shaped the community by controlling its structure from both the top-down and bottom-up, as discussed in the following sections.

### Top-down controls

#### Predatory/algicidal bacteria

Based on our phylogenetic trees (**Figure S8**), the most abundant ASV related to Saprospiraceae (ASV0037) was found to be most closely related to *Aureispira maritima. Aureispira*-related strains have been found to exhibit algicidal and predatory activity^45^, making ASV0037 a potential top-down controller of the Johor Strait microbial community. In general, many of the members of the family Saprospiraceae were found to have algicidal or predatory properties. Due to this, some studies have suggested to use *Saprospira*-like organisms to control cyanobacteria blooms, but this has never been experimentally shown nor attempted.^45–47^

#### Viruses

One other source of top-down control are viruses. What viruses lack in biomass, they make up for in abundance^48^. As viruses lack their own machinery for reproduction, they hijack bacterial cells and reproductive abilities for their replication. This intrusion releases a confetti of new virions, various cellular parts, and dissolved organic matter in the process. In this way, viruses relate the top-down controls of a community by predation to the bottom-up controls by way of nutrient availability: marine phages recycle an estimated 20% of all microorganisms every day, therefore making them a major driver in ocean biogeochemistry, and thus its biology and community structure^49^.

The positive correlation between virus counts and bacterial abundances (**Figure 1, Figure 3**), as well as the fact that no algal blooms nor any microbial abundance that is out of proportions were observed, supports the well-known Kill-The-Winner hypothesis. In this model of phage top-down control of microbial communities, more viruses are infecting the fastest-growing strain of microbes, killing the potential winner^50^. No one species is allowed to dominate the community, which allows other microbes to persist and thus increases the diversity and resilience of the system. This method of top-down control may be the primary mechanism of keeping the community structure in check.

### Bottom-up controls

#### Limiting nutrient availability

Chlorophyll measurements are a proxy for microbial biomass, especially photosynthetic microbes. We saw strong positive correlations between chlorophyll and N ratios (N:P and N:Si) that were consistent across all sites (**Figure 1**). As the increase in N compared to P or Si also results in an increase in microbial biomass, we suggest that nitrogen is the limiting nutrient in preventing the over-proliferation of photosynthetic microorganisms. This has also been mentioned in past studies^10, 12^.

#### Harmful Algal Blooms may be controlled from both the top-down and the bottom-up

Despite the fact that no HABs were observed during the sampling period, past HAB species were observed in 81 out of 118 samples in our time series. Similar to the more common taxa alternating more rapidly in the Johor Strait, these HAB taxa occur in very low numbers. However, the latter were not observed to dominate during the course of our sampling. Thus, we propose that an HAB event may develop only when two specific conditions are met: (1) an open niche is available and (2) top-down controls are no longer controlling the HAB population.

Two lines of evidence support the above hypothesis. Firstly, the results show that despite rapid and significant changes in the environmental conditions, the overall community structure recovers rapidly towards a stable centre-point. Moreover, changes in environmental parameters alone do not cause the system to crash and turn into an HAB-dominated community, showing the community’s resistance to disturbances. Secondly, there is strong evidence for the community to be controlled from the higher trophic levels by predation and viral lytic activities. While we do not have data on larger predators such as copepods and other zooplankton, viruses show changes that are synchronized with the variations in the bacterial populations (**Figure 3**).

The dominant species in the Johor Strait microbial community was shown to alternate between a few species depending on the nutrient conditions of the water. Changing environmental factors, coupled with the fact that HAB species were nearly always present in low amounts in the water, support the hypothesis that an HAB event may happen due to an available, open niche. A 2019 Singaporean study had proposed something similar: an observed dinoflagellate *Karenia* bloom may have happened due to a niche being opened whereby the abundance of diatoms had previously decreased^13^.

While the system seems to be able to adapt to changes in nutrient availability (bottom-up controls), it does not show adaptations to changes from the top-down (predators). This may be because top-down controls are more reactive in nature than causative; top-down controls do not initiate changes in the organisms at lower trophic levels, rather, top-down controls respond to the varying abundance of organisms at the lower trophic levels. When bottom-up controls make way for an open niche, HABs may be prevented from happening by the presence of predators limiting the uncontrolled growth of the HAB species. However, if their growth is somehow not inhibited by the top-down controls, the HAB species may take over and eventually dominate the microbial community, and thus an occurence of an HAB. This top-down control of algal blooms has been reported in several studies, be it the onset of blooms due to the decrease in grazer populations^51–54^ or the termination of blooms due to viral lytic activities^49, 55–58^. Irigoien et. al (2005)^54^ have also proposed the “loophole theory” in which blooms are formed when they are able to find a ‘loophole’ and escape the grazing control.

In the Johor Strait, the performance of these top-down controls may be affected by two mechanisms. As mentioned in the sections above, water mixing in the Johor Strait is primarily driven by tides (**Figure S4**). Previous studies have also concluded that blooms happen more frequently during neap tides when tidal mixing is at its minimum, due to the higher cell densities during a stable water column^10, 12^. However, we propose that stratification during minimum tidal mixing may also prevent the predators (grazers, viruses) from reaching their target preys. Sufficient tidal mixing would prevent the isolation of HAB species, thus exposing them to predators and inhibiting the HAB species’ proliferation when they find an open niche.

Secondly, ammonia (NH_4_) is traditionally known to be toxic for aquatic grazers such as copepods and other zooplanktons^59–61^. When NH_4_ concentrations go above a certain threshold, grazers perish from the toxicity. NH_4_ may enter water bodies through runoff from land, sewage, and industrial water discharge, and the Johor Strait is bounded by two highly urbanised regions which may act as sources of nutrient input^3–7^.

These two mechanisms controlling predators (ex. grazers, viruses, or algicidal bacteria), as well as changes in nutrient conditions opening up new niches mentioned earlier, may strongly contribute to the occurrence of HABs in the Johor Strait.

### Limitations & Further Study

Sampling was conducted every other day to stretch resources to cover a sampling period of 2 months. This means that any changes with a resolution of less than 2 days could not be observed, thus limiting our study. Furthermore, we have been focusing on compositional resilience instead of functional ones as we were not able to distinguish dormant microbes and genes from active ones through 16S rRNA gene amplicons. Future work could focus on the functional resilience of the Johor Strait microbes: it will be interesting to see which genes are activated at different conditions in response to the disturbances, and whether these genes mediated the effects of the perturbations.

## CONCLUSION

Frequent disturbances have shaped the Johor Strait microbial community by selection of, adaptation of, and diversification of organisms that can withstand these perturbations. The result is a diverse and stable community, adapted to living in a constantly changing environment. The increased importance of top-down controls in a highly productive environment such as this is in accordance with the Oksanen–Fretwell theory^62^, where the growth of bloom organisms is limited by predatory bacteria and viruses. Blooms may happen when an extreme disturbance is introduced into the system and changes both bottom-up or top-down controls, i.e., when the available empty niche temporally coincides with the demise of grazers or viral predators that keep the bloom populations in check. While we were not able to completely distinguish between the different disturbances, this study further highlights the complex interaction between perturbations and its role on regulating microbial community structure. Further studies may be focused on the functional regulation of the community, to see which active genes are expressed by the microorganisms in response to the constantly changing environment.

## ACKNOWLEDGMENTS

We thank SGUnited Trainee Ms Woo Yi Hui for assistance in sampling and DNA extraction & amplification, Dr. Syazwani Itri Amran (Universiti Teknologi Malaysia) for lending storage space in their -80°C freezer and liquid nitrogen supply during the pandemic, Dr. Cheng Dan and Dr. Mats Leifels (SCELSE) for guidance in flow-cytometry and analysis, Dr. Yongli Zhou for assistance in chlorophyll extraction and measurement, and finally Asst. Prof. Patrick Martin and Ms Oon Yee Woo (ASE) for their help in dissolved inorganic nutrient measurement and analysis. This project was funded by a collaboration grant from Center for Southeast Asian Coastal Interactions (SEACoast, CG3-HAB), as part of a broader research initiative supported by the Henry Luce Foundation. Winona Wijaya is supported by the National Research Foundation, Prime Minister’s Office, Singapore under its Competitive Research Programme (Award CRP21-2018-0005). Sandra Kolundzija is supported by a Singapore International Graduate Award (SINGA) from the Agency for Science, Technology & Research (A*STAR). We thank the committee members who organised the Joint Academic Microbiology Seminars (JAMS) 10th Annual Symposium, and JAMS for partially covering the publication costs for this article as part of the JAMS10 special issue in ISME communications. Planning and conducting field/lab work, and juggling various life stuff during a global pandemic is no joke, and we thank all our friends and families for loving and supporting us through all this.

## AUTHOR CONTRIBUTIONS STATEMENT

ZS and FML conceived the study. WW, ZS, CC, RS, SK, AMR, and NMA collected and processed the samples. WW, ZS, CC, and RS analyzed the data. WW, NHHC, and FML interpreted the results. WW drafted the manuscript. All authors edited the final version of the paper.

## ADDITIONAL INFORMATION

### Data availability

Raw sequencing data have been deposited to SRA under Bioproject number PRJNA848014. Processed data and scripts are available on https://github.com/winanonanona/2020-Time-Series.

## REFERENCES

1. Shade, A., Peter, H., Allison, S. D., Baho, D. L., Berga, M., Bürgmann, H. et al. Fundamentals of microbial community resistance and resilience. Front. Microbiol. 3, DOI: 10.3389/fmicb.2012.00417 (2012).

2. Philippot, L., Griffiths, B. S. & Langenheder, S. Microbial community resilience across ecosystems and multiple disturbances. Microbiol. Mol. Biol. Rev. 85, e00026–20, DOI: 10.1128/MMBR.00026-20 (2021).

3. Gin, K. Y.-H. Dynamics and size structure of phytoplankton in the coastal waters of Singapore. J. Plankton Res. 22, 1465–1484, DOI: 10.1093/plankt/22.8.1465 (2000).

4. Tan, K. S., Acerbi, E. & Lauro, F. M. Marine habitats and biodiversity of singapore’s coastal waters: A review. Reg. Stud. Mar. Sci. 8, 340–352, DOI: https://doi.org/10.1016/j.rsma.2016.01.008 (2016). Special Issue on the World Harbour Project — Global harbours and ports: different locations, similar problems?

5. Chai, X., Li, X., Hii, K. S., Zhang, Q., Deng, Q., Wan, L. et al. Blooms of diatom and dinoflagellate associated with nutrient imbalance driven by cycling of nitrogen and phosphorus in anaerobic sediments in Johor Strait (Malaysia). Mar. Environ. Res. 169, 105398, DOI: 10.1016/j.marenvres.2021.105398 (2021).

6. Goh, S. G., Bayen, S., Burger, D., Kelly, B. C., Han, P., Babovic, V. et al. Occurrence and distribution of bacteria indicators, chemical tracers and pathogenic vibrios in singapore coastal waters. Mar. Pollut. Bull. 114, 627–634, DOI: 10.1016/j.marpolbul.2016.09.036 (2017).

7. Hangzo, P. & Cook, A. D. The rise of iskandar malaysia: Implications for singapore’s marine and coastal environment. RSIS NTS Insight. IN14-01. DOI: 10.2139/ssrn.2392135 (2014).

8. Chénard, C., Wijaya, W., Vaulot, D., Lopes dos Santos, A., Martin, P., Kaur, A. et al. Temporal and spatial dynamics of Bacteria, Archaea and protists in equatorial coastal waters. Sci. Reports 9, 16390, DOI: 10.1038/s41598-019-52648-x (2019).

9. Tham, A. K. Seasonal distribution of the plankton in Singapore Straits. Mar. Biol. Assoc. India. 60–73. (May 1973)

10. Ooi, B. H., Zheng, H., Yue, K. P., Kurniawati, H., Sundarambal, P., Dao, M. H. et al. Case study of phytoplankton blooms in serangoon harbor of Singapore. In OCEANS’10 IEEE SYDNEY, 1–6, DOI: 10.1109/OCEANSSYD.2010.5603611 (IEEE, Sydney, Australia, 2010).

11. Leong, S. C. Y., Tkalich, P. & Patrikalakis, N. M. Monitoring harmful algal blooms in Singapore: Developing a HABs observing system. In 2012 Oceans - Yeosu, 1–5, DOI: 10.1109/OCEANS-Yeosu.2012.6263428 (IEEE, Yeosu, Korea (South), 2012).

12. Leong, S., Yew, C., Peng, L. L., Moon, C. S., Kit, J. K. W. & Ming, T. L. Three new records of dinoflagellates in Singapore’s coastal waters, with observations on environmental conditions associated with microalgal growth in the Johor Straits. RAFFLES BULLETIN OF ZOOLOGY 13 (2015).

13. Kok, J. W. K. & Leong, S. C. Y. Nutrient conditions and the occurrence of a Karenia mikimotoi (Kareniaceae) bloom within East Johor Straits, Singapore. Reg. Stud. Mar. Sci. 27, 100514, DOI: 10.1016/j.rsma.2019.100514 (2019).

14. Mohd-Din, M., Abdul-Wahab, M. F., Mohamad, S. E., Jamaluddin, H., Shahir, S., Ibrahim, Z. et al. Prolonged high biomass diatom blooms induced formation of hypoxic-anoxic zones in the inner part of Johor Strait. Environ. Sci. Pollut. Res. 27, 42948–42959, DOI: 10.1007/s11356-020-10184-6 (2020).

15. Lauro, F. M., Senstius, S. J., Cullen, J., Neches, R. Jensen, R. M., Brown, M. V. et al. The Common Oceanographer: Crowdsourcing the Collection of Oceanographic Data. PLoS Biol. 12, e1001947, DOI: 10.1371/journal.pbio.1001947 (2014).

16. Brussaard, C. P. D. Optimization of Procedures for Counting Viruses by Flow Cytometry. Appl. Environ. Microbiol. 70, 1506–1513, DOI: 10.1128/AEM.70.3.1506-1513.2004 (2004).

17. Brussaard, C. P. D., Payet, J. P., Winter, C. & Weinbauer, M. G. Quantification of aquatic viruses by flow cytometry. In Wilhelm, S., Weinbauer, M. & Suttle, C. (eds.) Manual of Aquatic Viral Ecology, 102–109, DOI: 10.4319/mave.2010.978-0-9845591-0-7.102 (American Society of Limnology and Oceanography, 2010).

18. R Core Team. R: A Language and Environment for Statistical Computing. R Foundation for Statistical Computing, Vienna, Austria (2021).

19. Wilkins, D., van Sebille, E., Rintoul, S. R., Lauro, F. M. & Cavicchioli, R. Advection shapes southern ocean microbial assemblages independent of distance and environment effects. Nat. Commun. 4, 2457, DOI: 10.1038/ncomms3457 (2013).

20. Allen, M. A. & Cavicchioli, R. Microbial communities of aquatic environments on heard island characterized by pyrotag sequencing and environmental data. Sci. Rep. 7, 44480, DOI: 10.1038/srep44480 (2017).

21. McNichol, J., Berube, P. M., Biller, S. J. & Fuhrman, J. A. Evaluating and improving small subunit rRNA PCR primer coverage for bacteria, archaea, and eukaryotes using metagenomes from global ocean surveys. mSystems. 6, e00565–21, DOI: 10.1128/mSystems.00565-21 (2021).

22. Martin, M. Cutadapt removes adapter sequences from high-throughput sequencing reads. EMBnet.journal. 17, 10–12, DOI: https://doi.org/10.14806/ej.17.1.200 (2011).

23. Callahan, B. J., McMurdie, P. J., Rosen, M. J., Han, A. W., Johnson, A. J. A. & Holmes, S. P. DADA2: High-resolution sample inference from illumina amplicon data. Nat. Methods. 13, 581–583, DOI: 10.1038/nmeth.3869 (2016).

24. Quast, C., Pruesse, E., Yilmaz, P., Gerken, J., Schweer, T., Yarza, P. et al. The SILVA ribosomal RNA gene database project: improved data processing and web-based tools. Nucleic Acids Res. 41, D590–D596, DOI: 10.1093/nar/gks1219 (2013).

25. Guillou, L., Bachar, D., Bittner, L., Boutte, C., Decelle, J., Edvardsen, B. et al. The protist ribosomal reference database (PR2): a catalog of unicellular eukaryote small sub-unit rRNA sequences with curated taxonomy. Nucleic Acids Res. 41, D597–D604, DOI: 10.1093/nar/gks1160 (2013).

26. Oksanen, J., Blanchet, F. G., Friendly, M., Kindt, R., Legendre, P., McGlinn, D. et al. vegan: Community ecology package. r package version 2.5-7.

27. McMurdie, P. J. & Holmes, S. phyloseq: An r package for reproducible interactive analysis and graphics of microbiome census data. PLoS One. 8, e61217, DOI: 10.1371/journal.pone.0061217.

28. Wickham, H. ggplot2: Elegant Graphics for Data Analysis (Springer-Verlag New York, 2016).

29. Kassambara, A. ggpubr: ‘ggplot2’ Based Publication Ready Plots (2020). R package version 0.4.0.

30. Kurtz, Z. D., Müller, C. L., Miraldi, E. R., Littman, D. R., Blaser, M. J. & Bonneau, R. Sparse and compositionally robust inference of microbial ecological networks. PLoS Comput. Biol. 11, e1004226, DOI: 10.1371/journal.pcbi.1004226.

31. Csardi, G. & Nepusz, T. The igraph software package for complex network research. InterJournal, Complex Systems 1695. 5, 1–9 (2006).

32. Bastian, M., Heymann, S. & Jacomy, M. Gephi: An open source software for exploring and manipulating networks. Proceedings of the International AAAI Conference on Web and Social Media 3, 361–362 (2009).

33. Wei, T. & Simko, V. R package ‘corrplot’: Visualization of a Correlation Matrix (2021). (Version 0.92).

34. O’Leary, N. A., Wright, M. W., Brister, J. R., Ciufo, S., Haddad, D., McVeigh, R. et al. Reference sequence (RefSeq) database at NCBI: current status, taxonomic expansion, and functional annotation. Nucleic Acids Res. 44, D733–D745, DOI: 10.1093/nar/gkv1189 (2016).

35. Katoh, K., Asimenos, G. & Toh, H. Multiple Alignment of DNA Sequences with MAFFT. Methods Mol. Biol. 537, 39–64, DOI: 10.1007/978-1-59745-251-9_3 (2009).

36. Price, M. N., Dehal, P. S. & Arkin, A. P. FastTree: Computing Large Minimum Evolution Trees with Profiles instead of a Distance Matrix. Mol. Biol. Evol. 26, 1641–1650, DOI: 10.1093/molbev/msp077 (2009). https://academic.oup.com/mbe/article-pdf/26/7/1641/13642970/msp077.pdf.

37. Suriyanti, S. & Usup, G. First report of the toxigenic Nitzschia navis-varingica (Bacillariophyceae) isolated from Tebrau Straits, Johor, Malaysia. Toxicon 108, 257–263, DOI: 10.1016/j.toxicon.2015.10.017 (2015).

38. Faust, K. Open challenges for microbial network construction and analysis. The ISME J. 15, 3111–3118, DOI: 10.1038/s41396-021-01027-4 (2021).

39. Röttjers, L. & Faust, K. From hairballs to hypotheses–biological insights from microbial networks. FEMS Microbiol. Rev. 42, 761–780, DOI: 10.1093/femsre/fuy030 (2018). https://academic.oup.com/femsre/article-pdf/42/6/761/26151423/fuy030.pdf.

40. Fuhrman, J. A., Cram, J. A. & Needham, D. M. Marine microbial community dynamics and their ecological interpretation. Nat. Rev. Microbiol. 13, 133–146, DOI: 10.1038/nrmicro3417 (2015).

41. Banerjee, S., Schlaeppi, K. & van der Heijden, M. G. A. Keystone taxa as drivers of microbiome structure and functioning. Nat. Rev. Microbiol. 16, 567–576, DOI: 10.1038/s41579-018-0024-1.

42. Hosoya, S., Arunpairojana, V., Suwannachart, C., Kanjana-Opas, A. & Yokota, A. Aureispira maritima sp. nov., isolated from marine barnacle debris. Int. J. Syst. Evol. Microbiol. 57, 1948–1951, DOI: 10.1099/ijs.0.64928-0 (2007).

43. Jacquet, C. & Altermatt, F. The ghost of disturbance past: long-term effects of pulse disturbances on community biomass and composition. Proc. R. Soc. B. 287, 20200678, DOI: 10.1098/rspb.2020.0678 (2020).

44. Renes, S. E., Sjötedt, J., Fetzer, I. & Langenheder, S. Disturbance history can increase functional stability in the face of both repeated disturbances of the same type and novel disturbances. Sci. Rep. 10, 11333, DOI: 10.1038/s41598-020-68104-0 (2020).

45. McIlroy, S. J. & Nielsen, P. H. The Family Saprospiraceae, In The Prokaryotes: Other Major Lineages of Bacteria and The Archaea. 863–889. DOI: 10.1007/978-3-642-38954-2_138 (Springer Berlin Heidelberg, Berlin, Heidelberg, 2014).

46. Ashton, P. J. & Robarts, R. D. Apparent predation of Microcystis aeruginosa kütz. emend elenkin by a Saprospira-like bacterium in a hypertrophic lake (Hartbeespoort Dam, South Africa). J. Limnol. Soc. South. Afr. 13, 44–47, DOI: 10.1080/03779688.1987.9634542 (1987).

47. Kang, Y.-H., Jung, S. W., Jo, S.-H. & Han, M.-S. Field assessment of the potential of algicidal bacteria against diatom blooms. Biocontrol Sci. Technol. 21, 969–984, DOI: 10.1080/09583157.2011.591922 (2011).

48. Kolundžija, S., Cheng, D.-Q. & Lauro, F. M. RNA viruses in aquatic ecosystems through the lens of ecological genomics and transcriptomics. Viruses. 14, 702, DOI: 10.3390/v14040702 (2022).

49. Suttle, C. A. Marine viruses — major players in the global ecosystem. Nat. Rev. Microbiol. 5, 801–812, DOI:10.1038/nrmicro1750 (2007).

50. Thingstad, T. F. Elements of a theory for the mechanisms controlling abundance, diversity, and biogeochemical role of lytic bacterial viruses in aquatic systems. Limnol. Oceanogr. 45, 1320–1328, DOI: 10.4319/lo.2000.45.6.1320 (2000).

51. Buskey, E. J., Montagna, P. A., Amos, A. F. & Whitledge, T. E. Disruption of grazer populations as a contributing factor to the initiation of the Texas brown tide algal bloom. Limnol. Oceanogr. 42, 1215–1222, DOI: 10.4319/lo.1997.42.5 part 2.1215 (1997).

52. Buskey, E. J. How does eutrophication affect the role of grazers in harmful algal bloom dynamics? Harmful Algae 8, 152–157, DOI: 10.1016/j.hal.2008.08.009 (2008).

53. Glibert, P. M., Berdalet, E., Burford, M. A., Pitcher, G. C. & Zhou, M. Harmful Algal Blooms and the Importance of Understanding Their Ecology and Oceanography. In Global Ecology and Oceanography of Harmful Algal Blooms, vol. 232, 9–25, DOI: 10.1007/978-3-319-70069-42 (Springer International Publishing, Cham, 2018). Series Title: Ecological Studies.

54. Irigoien, X., Flynn, K. J. & Harris, R. P. Phytoplankton blooms: a ‘loophole’ in microzooplankton grazing impact? J. Plankton Res. 27, 313–321, DOI: 10.1093/plankt/fbi011 (2005).

55. Lehahn, Y., Koren, I., Schatz, D., Frada, M., Sheyn, U., Boss, E. et al. Decoupling physical from biological processes to assess the impact of viruses on a mesoscale algal bloom. Curr. Biol. 24, 2041–2046, DOI: https://doi.org/10.1016/j.cub.2014.07.046 (2014).

56. Brussaard, C. P. D., Wilhelm, S. W., Thingstad, F., Weinbauer, M. G., Bratbak, G., Heldal, M. et al. Global-scale processes with a nanoscale drive: the role of marine viruses. The ISMEJ. 2, 575–578, DOI: 10.1038/ismej.2008.31 (2008).

57. Vincent, F., Sheyn, U., Porat, Z., Schatz, D. & Vardi, A. Visualizing active viral infection reveals diverse cell fates in synchronized algal bloom demise. Proc. Natl. Acad. Sci. 118, e2021586118, DOI: 10.1073/pnas.2021586118 (2021).

58. Wilhelm, S. W. & Suttle, C. A. Viruses and Nutrient Cycles in the Sea. 49, 8 (1999).

59. Arauzo, M. Harmful effects of un-ionised ammonia on the zooplankton community in a deep waste treatment pond. Water Res. 37, 1048–1054, DOI: 10.1016/S0043-1354(02)00454-2 (2003).

60. Arauzo, M. & Valladolid, M. Short-term harmful effects of unionised ammonia on natural populations of moina micrura and brachionus rubens in a deep waste treatment pond. Water Res. 37, 2547–2554, DOI: 10.1016/S0043-1354(03)00023-X (2003).

61. Sullivan, B. K. & Ritacco, P. Ammonia toxicity to larval copepods in eutrophic marine ecosystems: A comparison of results from bioassays and enclosed experimental ecosystems. Aquat. Toxicol. 7, 205–217, DOI: 10.1016/S0166-445X(85)80006-0 (1985).

62. Oksanen, L., Fretwell, S. D., Arruda, J., & Niemela, P. Exploitation Ecosystems in Gradients of Primary Productivity. The American Naturalist, 118, 240–261, (1981). http://www.jstor.org/stable/2460513

63. Hydrographic Department, Maritime and Port Authority of Singapore. Singapore Tide Tables Year 2020 (Singapore, 2019).

