## Supplementary File for "Frequent pulse disturbances influence resistance and resilience in tropical marine microbial communities"

**Table S1** Details of sampling sites

| Site | Coordinates | E/W | N/S | Time of sampling |
|---|---|---|---|---|
| Sembawang (SBW) | 1.4643, 103.8372 | East | South | 9.30am |
| Senibong (SNB) | 1.4846, 103.8170 | East | North | 9.30am |
| Woodlands (WDL) | 1.4554, 103.7790 | West | South | 11.00am |
| Stulang (STL) | 1.4665, 103.7791 | West | North | 11.00am |

**Table S2** Dates of spring and neap tides, and their related moon phases

| Moon Phase | Tide | Date (dd/mm/yyyy) |
|---|---|---|
| Full moon | Spring tide | 31/10/2020 |
| Last quarter | Neap tide | 08/11/2020 |
| New moon | Spring tide | 15/11/2020 |
| First quarter | Neap tide | 22/11/2020 |
| Full moon | Spring tide | 30/11/2020 |
| Last quarter | Neap tide | 08/12/2020 |
| New moon | Spring tide | 15/12/2020 |
| First quarter | Neap tide | 22/12/2020 |
| Full moon | Spring tide | 30/12/2020 |

**Table S3** Sembawang Tide station measures tides at 1.830m below the Singapore Height Datum, which is 1.652m above the Chart Datum[63]. Along the course of our sampling, the highest tide height measures at 3.74m, the lowest at 0.04m, and the average tide height is 2.08m. We are unable to publish the tide data, but this table shows the overview of historical tide heights at the Sembawang Tide Station.

| Sembawang Tide Station | Height (m) |
|---|---|
| Mean High Water Springs | 3.1 |
| Mean High Water Neaps | 2.5 |
| Mean Level | 1.9 |
| Mean Low Water Neaps | 1.4 |
| Mean Low Water Springs | 0.7 |

**Table S4** Summary of CCA results (**Figure S7**) of the top few taxa, i.e. the location on each arrow where a line perpendicular to the arrow and the ASV can be drawn.

| Taxa | ASV No. | DIN | Si | PO4 | DIN:PO4 | DIN:Si | Salinity |
|---|---|---|---|---|---|---|---|
| *(Genus)* HIMB11 | 3 | Low | Low | Low | Mid | High | Mid-high |
| *(Genus)* PCC-6307 | 4, 7 | High | High | High | Mid-low | Low | Mid-low |
| *(Genus)* Nitrosopumilus | 9, 16 | Mid | High | Mid-high | Low | Mid-low | High |
| *(Family)* Flavobacteriaceae | 13, 15 | Mid-low | Low | Low | Mid-high | High | Mid-high |
| *(Family)* Saprospiraceae | 37 | High | Low | Very low | Very high | Very high | Very low |

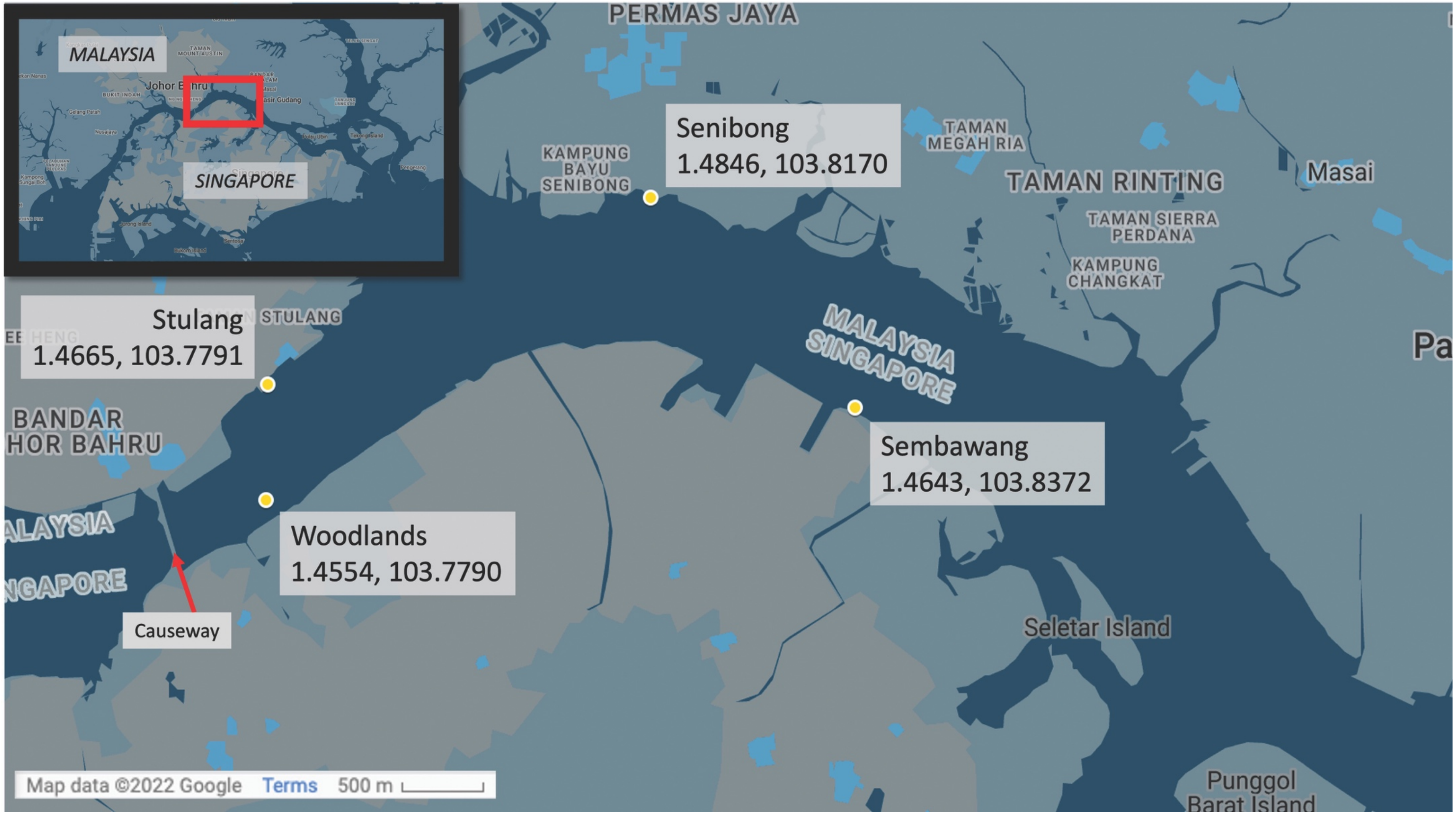


**Figure S1** Map of Singapore showing the four sampling stations in this study.

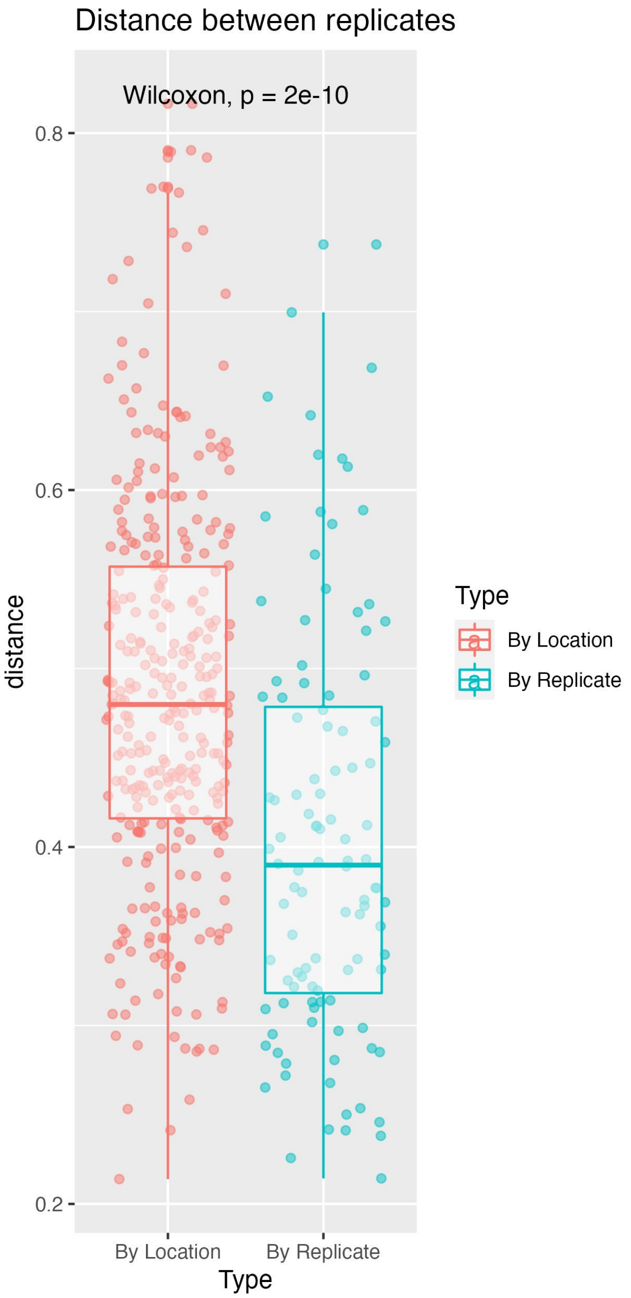


**Figure S2** Bray-Curtis distance between different locations on the same day and different technical replicates. The distance between technical replicates is smaller than between locations.

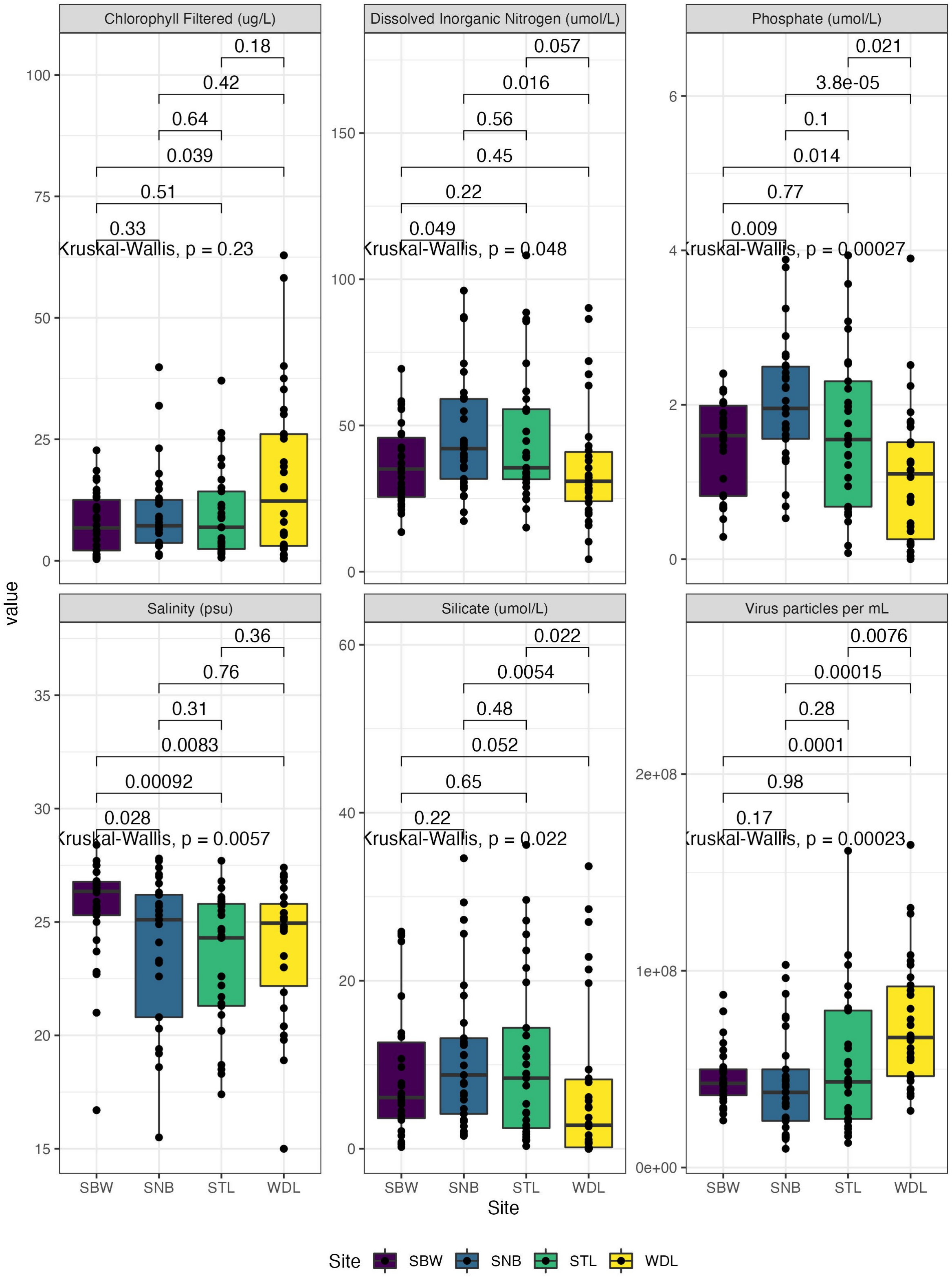


**Figure S3** Summary of dissolved nutrient values and environmental parameters. Using Bonferroni correction for multiple comparisons, only p-values below 0.00833 show significance.

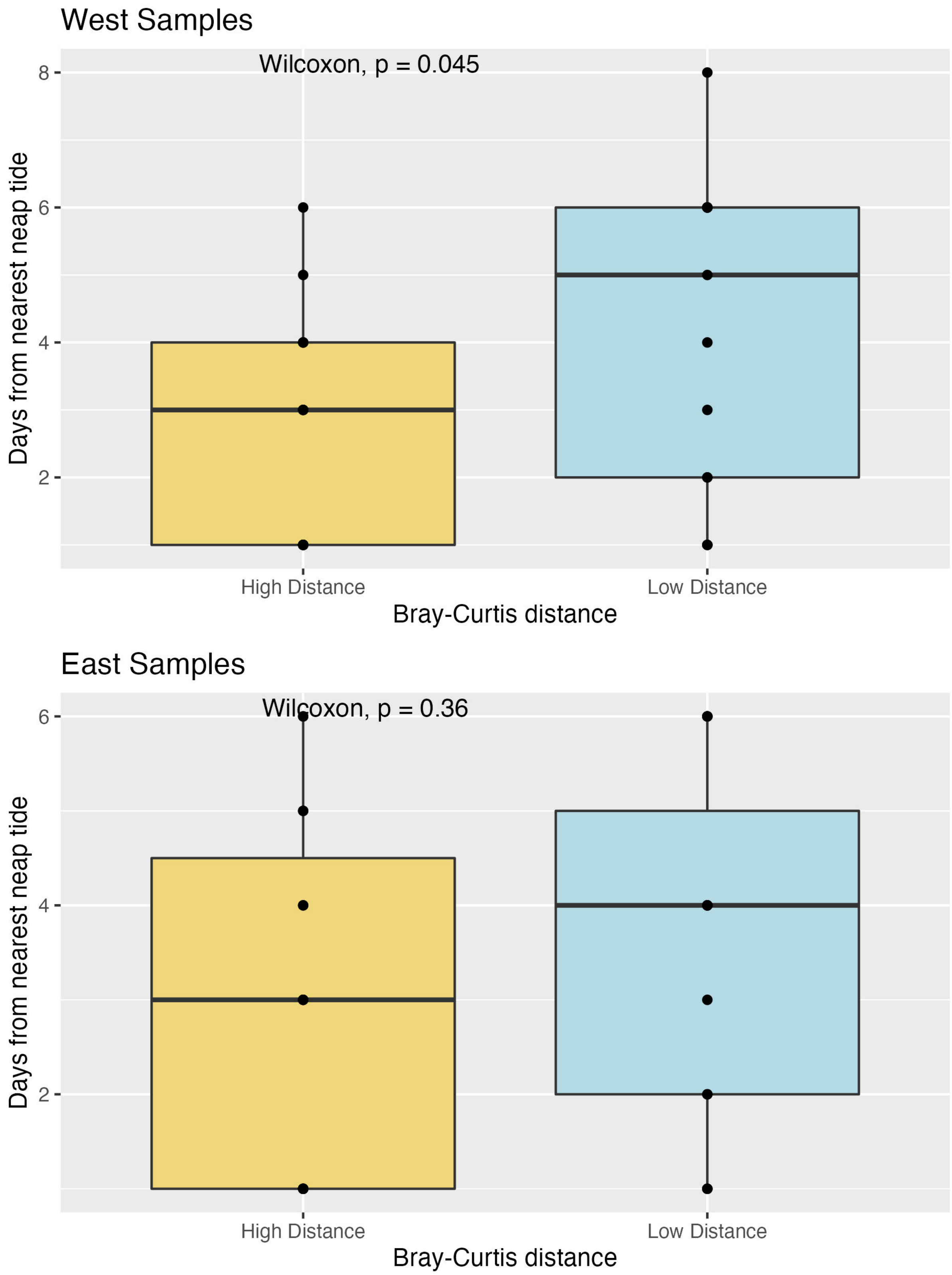


**Figure S4** Days from neap tides in samples with a high or low Bray-Curtis distance from the daily average. A neap tide happens when the moon is at a 90° angle with the earth and the sun, causing high tides to be higher and low tides to be lower, thus a lower tidal mixing. Further away from a neap tide, the water experiences higher tidal mixing. There is a significant difference between the high and low distance samples with regard to tidal mixing in the western samples (top), while the difference is insignificant in the eastern samples (bottom). This shows a higher influence of tidal-driven mixing in the western samples closer to the causeway, than in the eastern samples closer to the South China Sea.

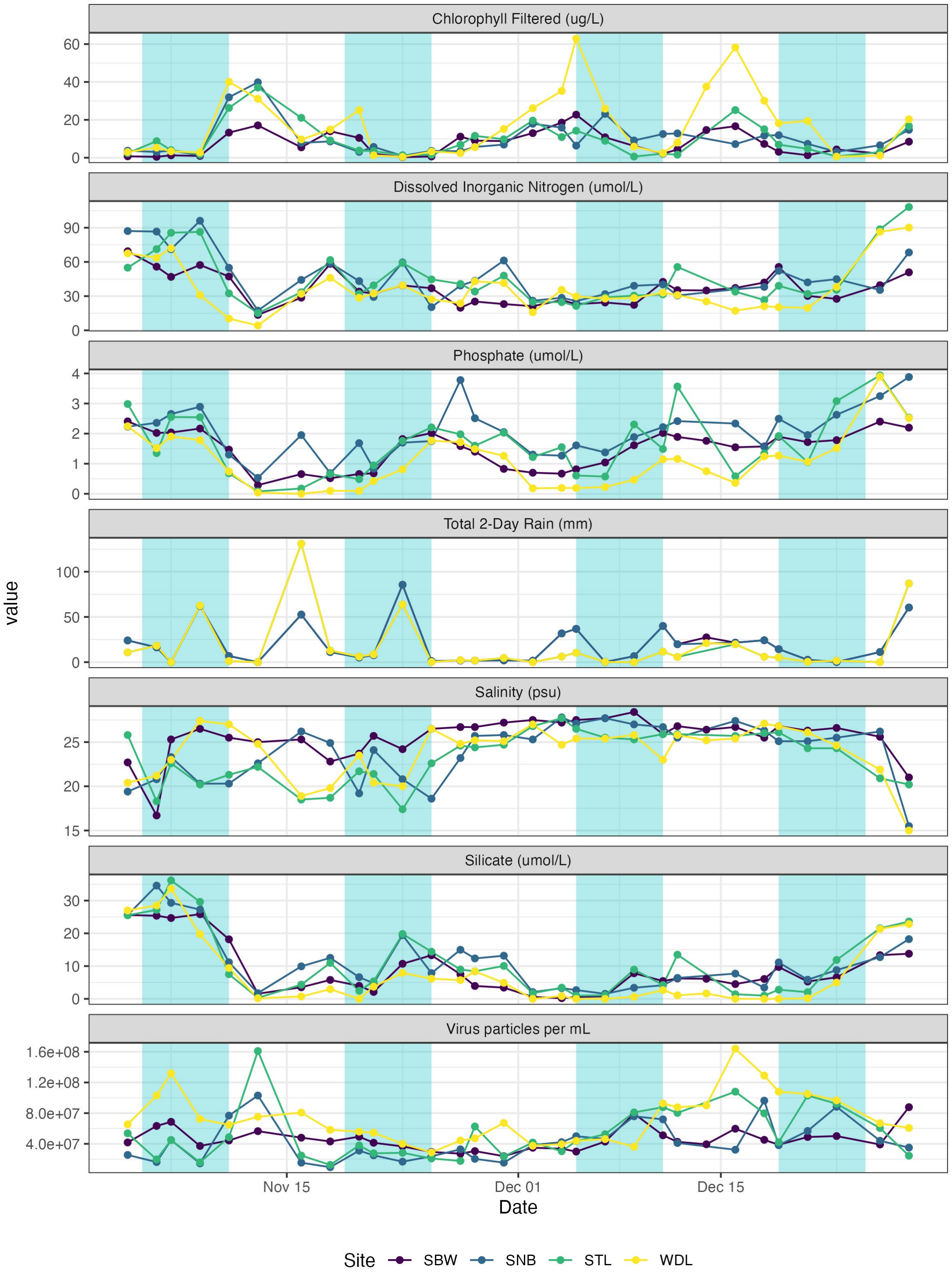


**Figure S5** A time-series of environmental parameters and nutrient values. 3 days before/after the peak of the neap tide is shaded blue.

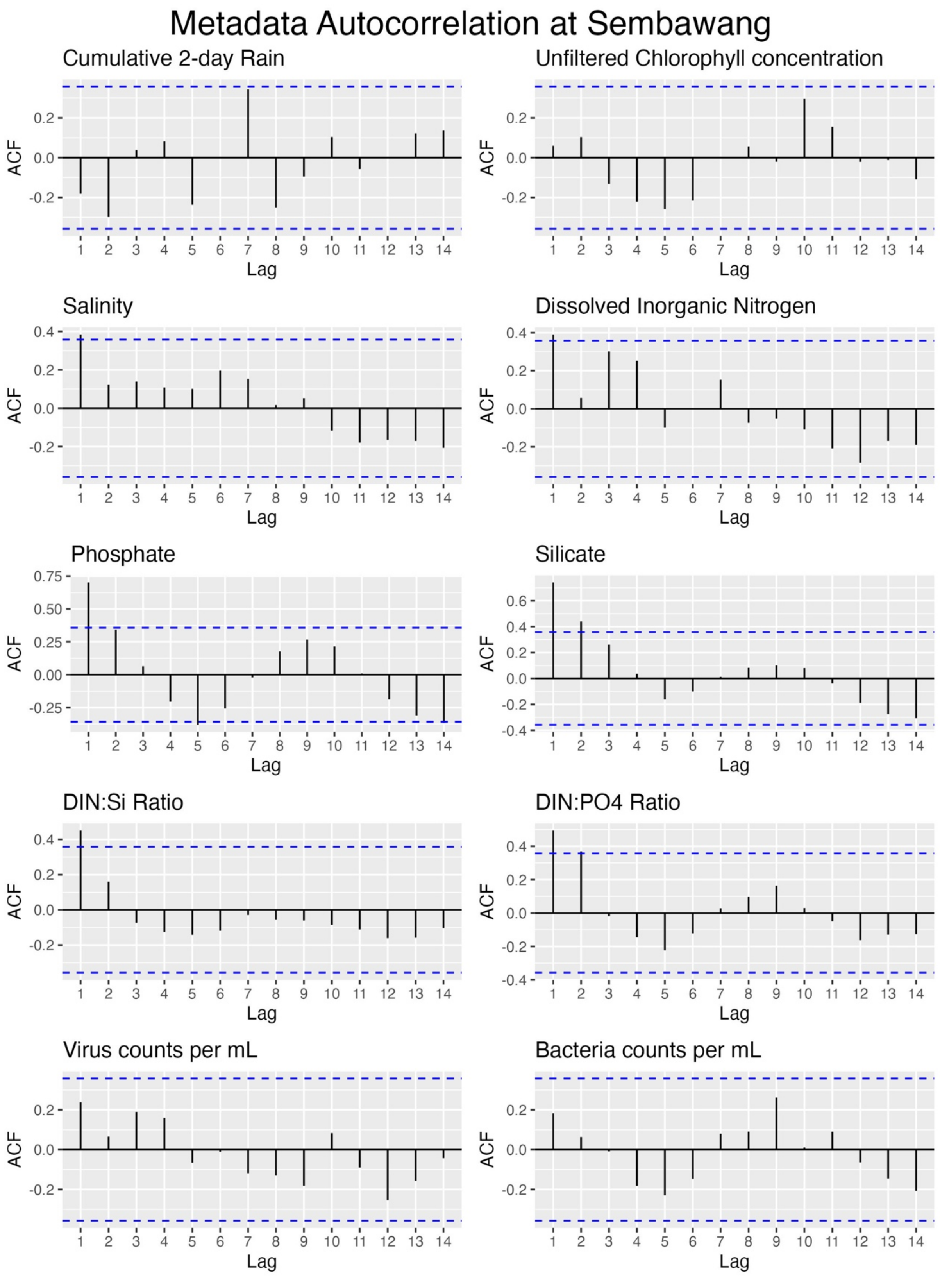


**Figure S6** Autocorrelation of metadata. Blue dotted line marks significant autocorrelation values. No significant autocorrelation was detected. Only autocorrelation of Sembawang metadata was shown; autocorrelation plots of Senibong, Stulang, and Woodlands show similar results.

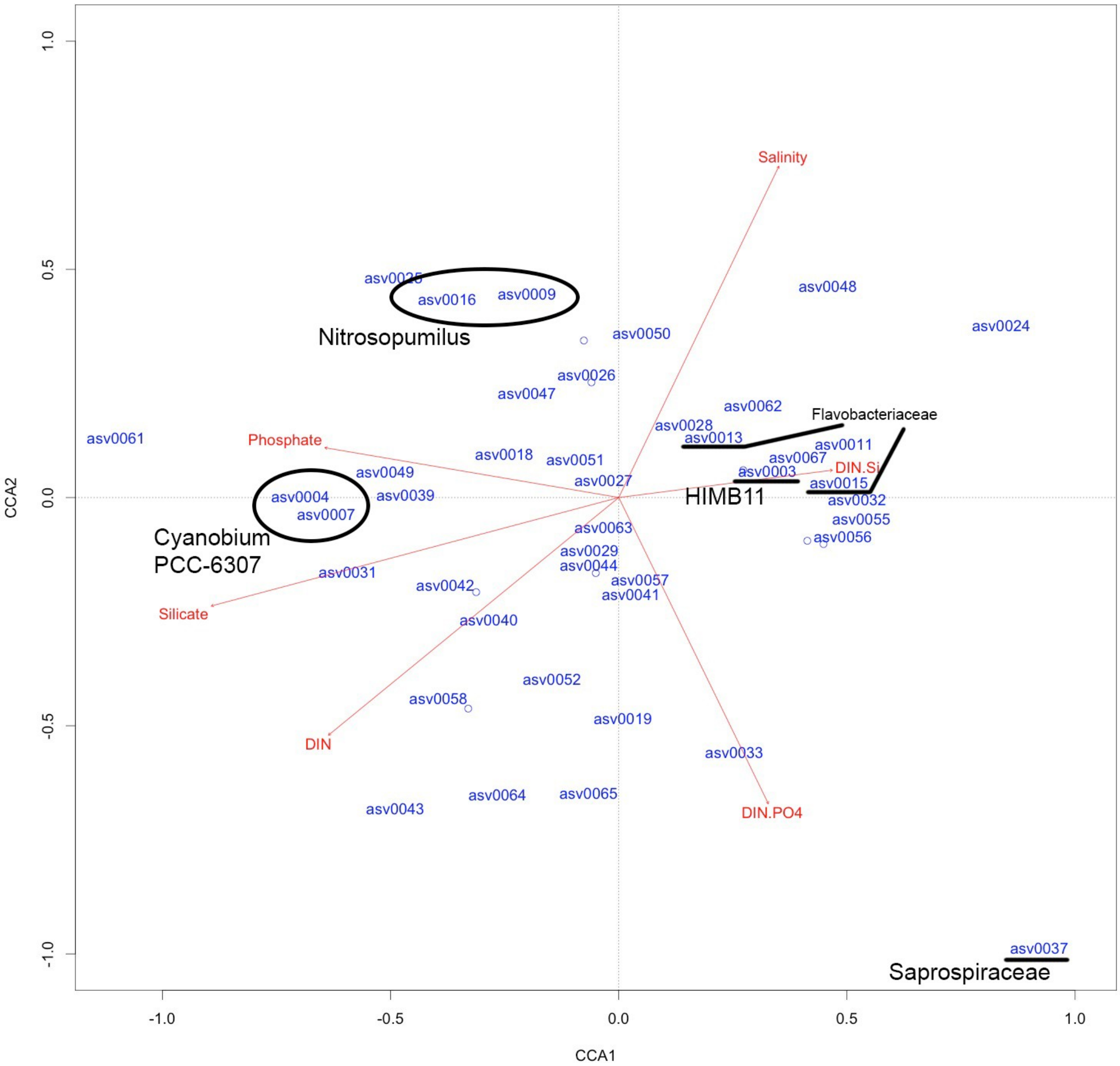


**Figure S7** Canonical Correspondence Analysis (CCA) of the top 50 ASVs. The arrows show various environmental parameters. A few notable ASVs has been annotated with their taxonomic classification. **Table S4** summarises each taxon's relationship to the environmental parameters, i.e. the location on each arrow where a line perpendicular to the arrow and the ASV can be drawn.

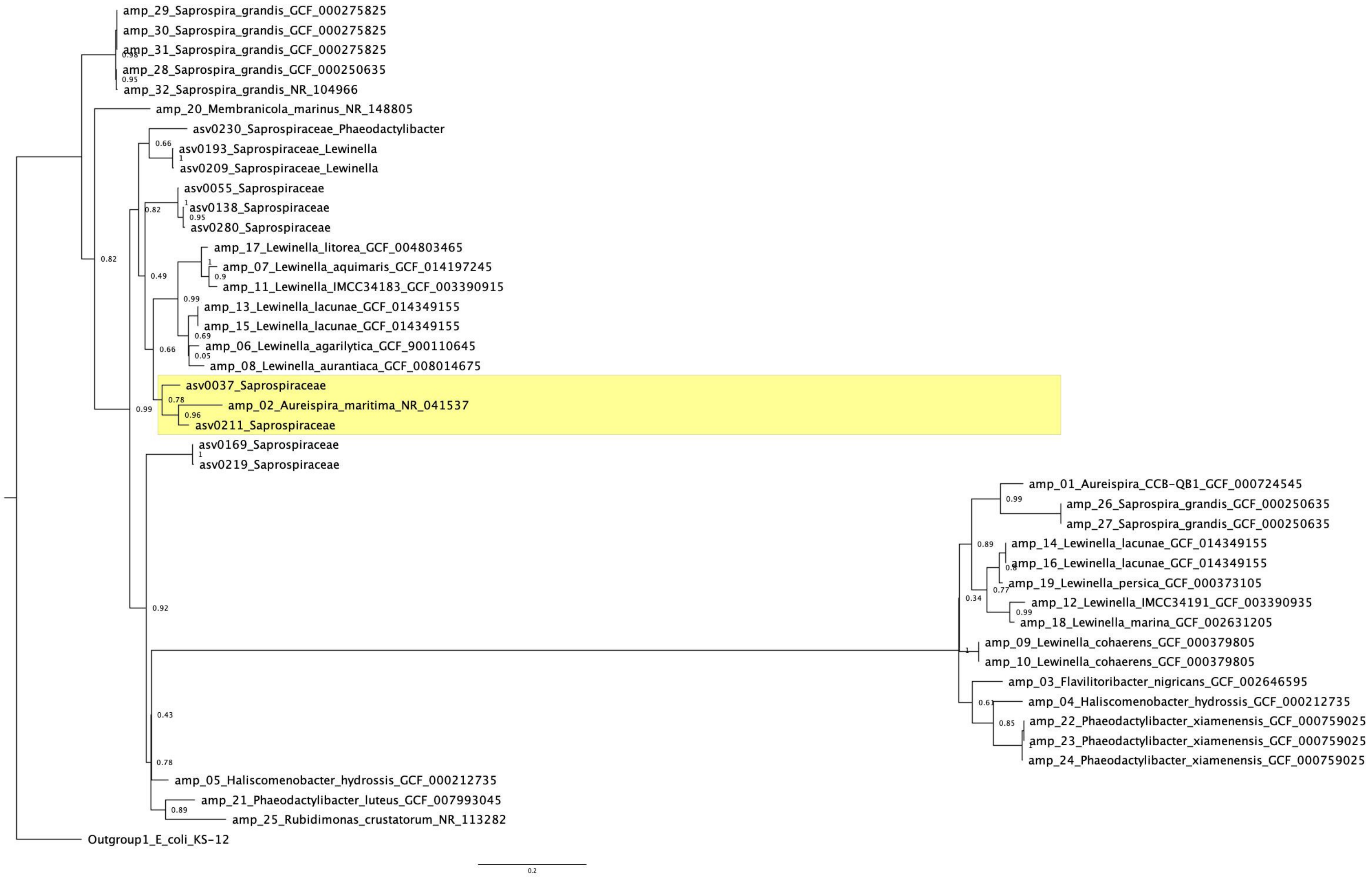


**Figure S8** A Maximum-likelihood phylogenetic tree showing Saprospiraceae-related ASVs and in silico amplicons taken from the NCBI RefSeq database. Highlighted features contain ASV0037, the most abundant Saprospiraceae ASV, as well as its potentially closest relative, *Aureispira maritima*. Numbers on each split show the local support values calculated using the Shimodaira-Hasegawa test with 1000 resamples[36]. Codes starting with "GCF" or "NR" are NCBI reference numbers of the genome.
